# Tempo and timing of ecological trait divergence in bird speciation

**DOI:** 10.1101/083253

**Authors:** Jay P. McEntee, Joseph A. Tobias, Catherine Sheard, J. Gordon Burleigh

## Abstract

Organismal traits may evolve either gradually or in rapid pulses followed by periods of stasis, but the relative importance of these evolutionary models in generating biodiversity has proven difficult to resolve^1,2^. In addition, while it is often assumed that pulses of trait evolution are associated with speciation events, few studies have explicitly examined how the tempo of trait divergence varies with respect to different geographical phases of speciation. Thus, we still know little about the trajectories of trait divergence over timescales relevant to speciation, or the extent to which these trajectories are shaped by variation in geographical isolation and overlap (sympatry) among incipient species. Here, we combine divergence time estimates, trait measurements, and geographic range data for avian sister species pairs worldwide to examine the tempo and timing of trait divergence during allopatric speciation. We show that divergence in two important ecological traits—?body mass and beak morphology—is best explained by a model including pulses of divergence and periods of relative stasis. We also infer that trait divergence pulses often precede sympatry, and that pulses leading to greater trait disparity are associated with earlier transitions to sympatry. These findings suggest that early pulses of trait divergence promote subsequent transitions to sympatry, rather than such pulses occurring after sympatry has been established, for example via character displacement^3^. Incorporating pulsed divergence models into allopatric speciation theory helps to resolve some apparently contradictory observations, including widespread instances of both rapid sympatry and prolonged geographical exclusion^4-6^.

## TEXT

Speciation in vertebrates may proceed over long and variable periods^7^. The entire process from onset to completion is often subdivided into three stages, beginning with a phase of geographic isolation (allopatry), followed by secondary contact, and finally the transition to coexistence in overlapping geographical ranges (sympatry; see Fig. 1)^7-10^. In standard forms of this model, the third stage is delayed by competitive interactions^11^ or incomplete reproductive isolation^9^, and thus only occurs when traits are sufficiently divergent to facilitate sympatry^6^. However, while the pattern of increased trait divergence in sympatric lineages is widespread among animal taxa^4^, the timing and geographical context of the process of trait divergence is often unclear. In particular, trait divergence could arise primarily by the accumulation of differences prior to secondary contact^12^ or alternatively after sympatry is established^3^. Furthermore, the tempo and mode of trait divergence during speciation is also controversial, with some studies describing divergence as slow or gradual throughout the process^5,12^, while others provide evidence for abrupt, pulse-like changes^13,14^ occurring either early in allopatry^15,16^ or later in sympatry^3, 6^ (Fig. 1). It can even be argued that ecological (local) adaptation in allopatry, followed by species interactions in sympatry, provide the context for multiple pulses of trait divergence over time^15^.

**Figure 1.**
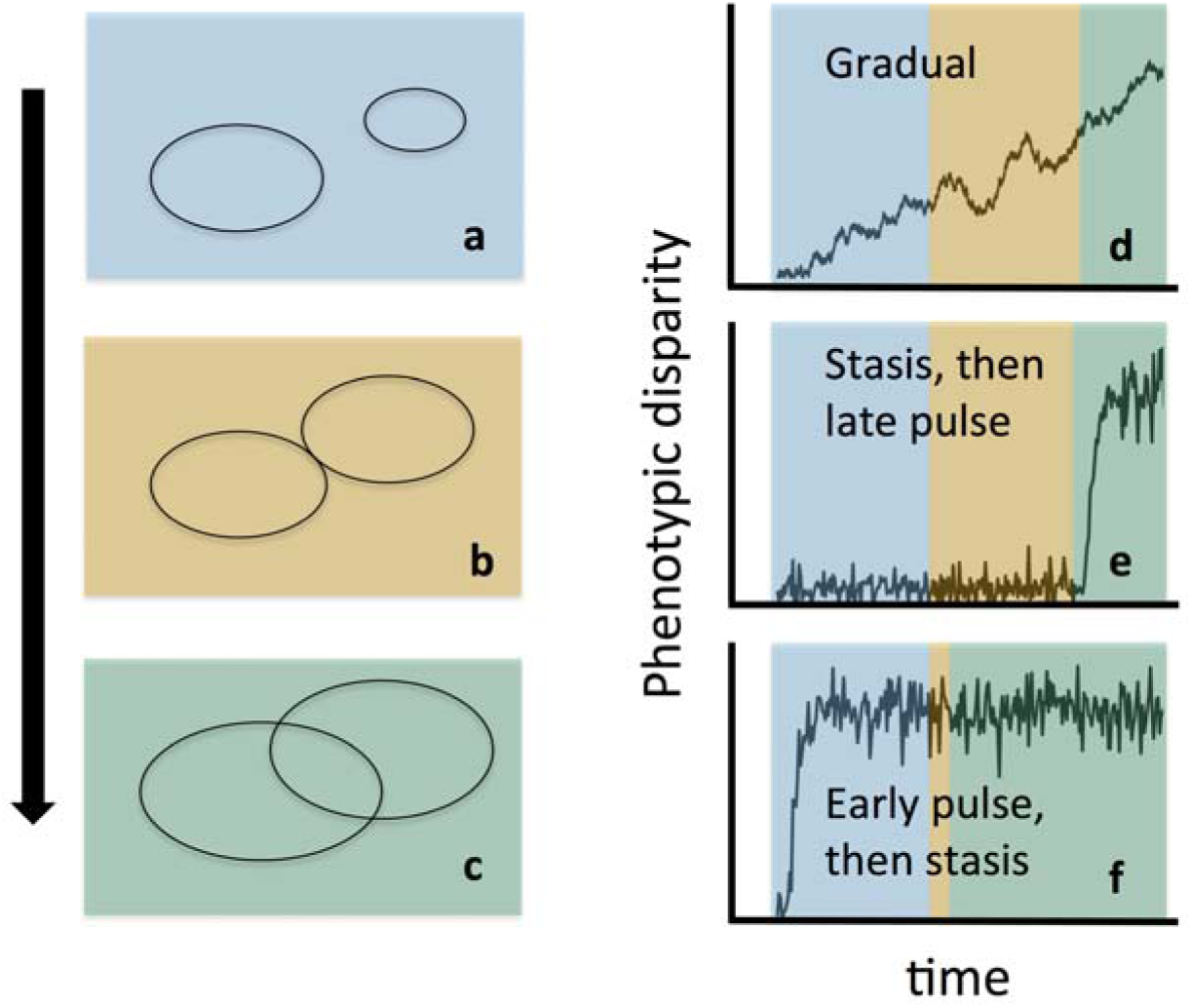
The speciation cycle and phenotypic trait divergence. Bird speciation typically involves a sequence of geographical states, starting with an allopatric phase (**a**), followed by secondary contact (**b**), and finally sympatry (**c**). Phenotypic divergence may take different pathways during this cycle: gradual models predict no pulse of divergence at any point in the cycle (**d**), whereas punctuated models involve stasis punctuated by pulses, which can follow the onset of coexistence (**e**) or precede it (**f**). Note that secondary contact (orange) is extended when traits are similar (**e**), and reduced when traits have already substantially diverged in allopatry (**f**).

Prolonged speciation creates problems when modeling pulsed trait evolution at a macroevolutionary scale, and hence for most studies comparing evidence for pulsed versus gradual evolution. Pulsed models typically assume that evolutionary change is concentrated at speciation^2^, which is modelled as a single instantaneous event, potentially leading to misinterpretations about trait evolution. Moreover, the contribution of pulsed trait evolution may become obscured because stasis (or bounded evolution^17^) with intermittent pulses can resemble gradualism at these coarse macroevolutionary scales^2^. Micro-evolutionary (population-level) studies conducted over short time periods confirm that stasis is prevalent^18^ and also that trait evolution pulses sometimes occur^4,17^. Observed pulse events, however, may represent brief departures of trait values from their long-term means, followed by reversion, and may therefore not contribute strongly to patterns of trait variation at macroevolutionary scales^4,17, 18^. Thus, although previous studies have found support for both gradual and pulsed modes of evolution^19,20^, their relative importance in generating biological diversity remains unknown^2^. An added complication is that trait divergence during the allopatric phase of speciation may be ephemeral if gene pools merge during secondary contact, whereas greater levels of divergence may lead to reproductive isolation and ultimately sympatry, a possibility that could accentuate patterns of pulsed evolution in phylogenetic approaches and the fossil record, even when divergence itself is gradual^16,21^.

Disentangling these alternative divergence pathways is a key step in resolving general patterns of trait evolution and predicting which nascent species ultimately succeed, leaving daughter species, and which will perish or fail to remain distinct when changing environments re-organize geographic ranges^16,21, 22^. However, our understanding of the rates and timing of trait divergence in relation to stages of the speciation process in vertebrates remains highly incomplete, not least because the data required to test these ideas are often lacking. In particular, although broad-scale information on ecological traits and phylogenetic history is available for some large vertebrate clades^20^, the accompanying information on geographic ranges is not sufficiently resolved to explore divergence pathways in the context of geographical phases of speciation.

To address this issue, we examined phenotypic divergence, geographic relationships, and estimated divergence times^23-25^ among 952 pairs of avian sister species. As we were interested in the tempo and timing of divergence during the speciation process, we applied a set of evolutionary models designed to span both microevolutionary and macroevolutionary processes^17^ to estimated trait disparities and divergence times. We used this approach because species pairs may vary in the extent to which their trait divergence is better characterized by microevolutionary or macroevolutionary processes. Focusing on two important ecological traits—body mass and beak morphology—we assessed relative support for four stochastic trait divergence models: “gradual”, “single pulse”, “multiple pulse”, and time-independent (“white noise”) models (see Methods). The first three models (i.e. all except the white noise model) incorporate a bounded evolution component to represent processes at shorter timescales. We found the strongest support for the single pulse model, in which the bounded evolution component is relatively narrow^26^ (Supplementary Tables 10-13). Support for the single pulse model was much stronger than for the gradual model (ΔAIC 899 for body mass, ΔAICs 862, 1004, and 968 for beak PC1, PC2, and PC3, respectively; see Methods and Supplementary Tables 10-13), in agreement with phylogenetic studies reporting a strong contribution of pulses in the accumulation of phenotypic diversity^19,27^.

Moreover, these results raise two further questions that we address here: 1) how does the estimated timing of divergence pulses compare to typical progressions through the geographic speciation process (allopatry, secondary contact, sympatry)? and 2) does pulsed ecological trait divergence impact transitions through this process? With respect to the first question, the expected waiting time to a pulse of divergence in the single pulse model was ∼670,000 years (95% CI from likelihood profile: 280,000 years to 1.13 My) for body mass (see also Figs. 3 and 4, Supplementary Table 10). The expected waiting times to a pulse in beak morphology divergence in single pulse models were ∼560,000 years for PC1 (95% CI: 200,000 years to 1.0 My), ∼170,000 years for PC2 (95% CI: 0 to 410,000 years), and ∼90,000 years for PC3 (95% CI: 0 to 280,000 years; see Supplementary Tables 11-13). To compare these estimates with progression to secondary contact and sympatry, we used a fine-grained geospatial database of ∼178 million species observation records and standard geographical range polygons, respectively (see Methods). Secondary contact often occurs so rapidly that the signature of allopatry is difficult to detect in our analyses of local co-occurrence (contact) and divergence time (Extended Data Fig. 4), suggesting that pulses may occur following secondary contact or even during parapatric speciation (see below, Methods and Supplementary Information). However, based on the relative timescales of trait divergence pulses and sympatry establishment shown in (Fig. 4), we also conclude that divergence pulses typically precede the establishment of sympatry, and are thus unlikely to be driven by character displacement processes^3^.

**Figure 3.**
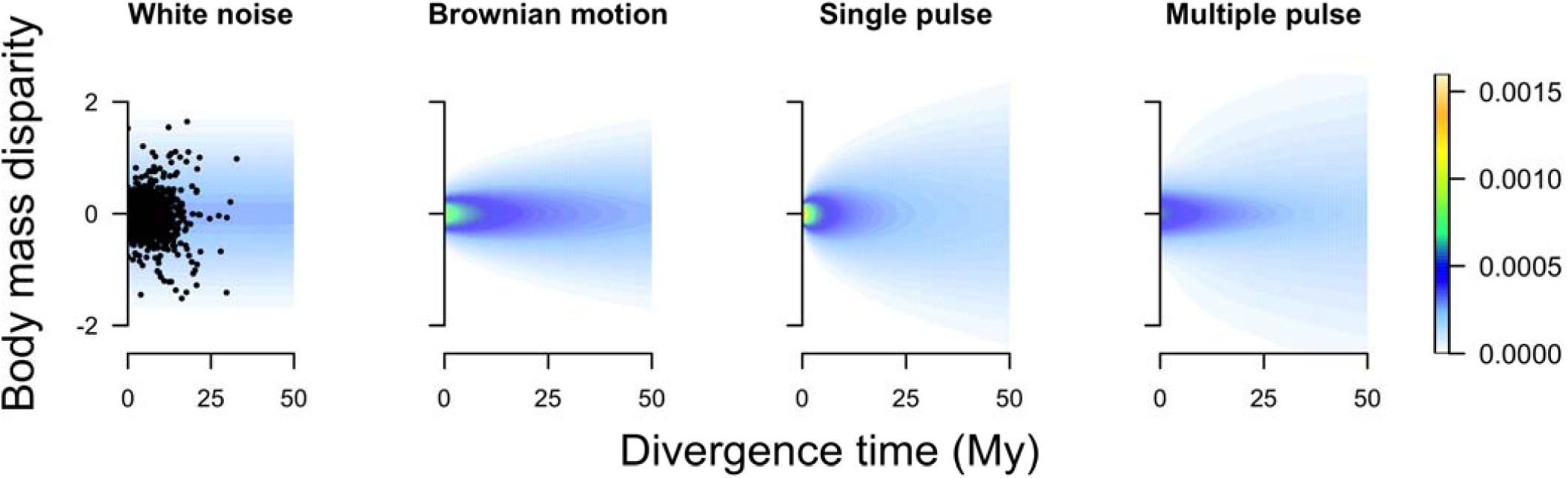
Tempo of body mass divergence for avian sister species. Stochastic pulsed models provide better fits to patterns of body mass divergence and divergence time, with the best fit a single pulse model (ΔAIC relative to the multiple pulse model: 797). Colors denote probability density. The probability density for any time slice follows a normal distribution (most apparent in the white noise model where the probability density distribution is independent of time). Relative probability density can be assessed within each time slice but not across time. For clarity, the empirical data points are plotted only on the white noise model.

**Figure 4.**
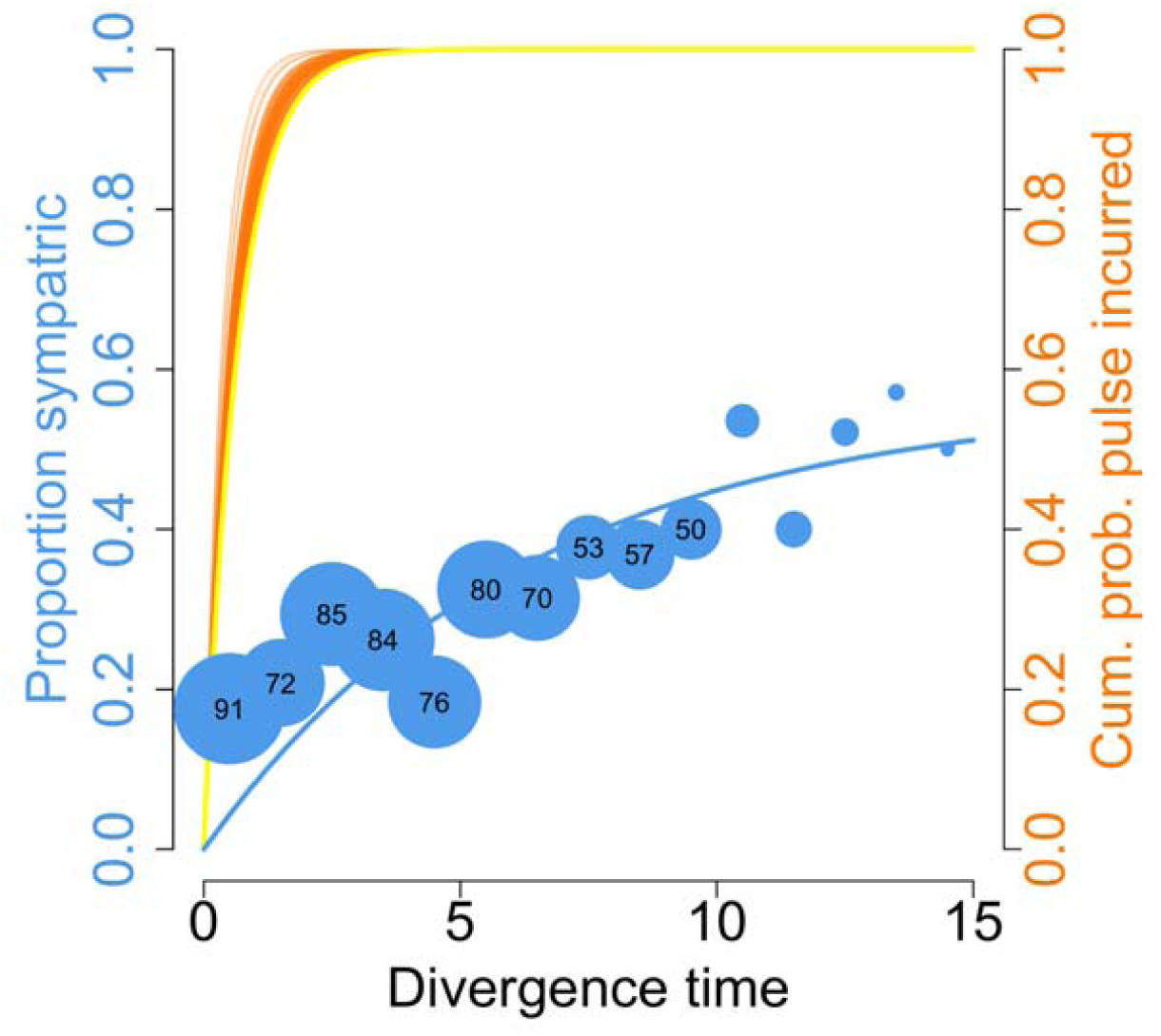
Timing of body mass divergence pulses and sympatry. Comparison of timescales suggests that mass divergence tends to precede sympatry among 952 avian sister species. Yellow and orange lines are cumulative probability distributions of incurring a pulse under the single pulse model (yellow: estimated from maximum likelihood phylogenetic tree^23^; orange: from 100 bootstrap trees^23^). Circles are proportions of sympatric species pairs for 1-million year intervals of divergence time^23^; circle sizes represent sample sizes, numbered where ≥50. For visual comparison, an exponential decay model has been fitted to the proportion of sister pairs in sympatry (blue curve; assumes sympatry is secondary).

Early pulses of ecological trait divergence theoretically reduce both competition and reproductive interference among incipient species^15^, potentially overcoming constraints on sympatry^10,22^. To assess whether such pulses influence rates of transition through geographical stages of the speciation process, we tested whether variation in body mass and beak morphology predicted which species pairs are parapatric or sympatric. Focusing on all species pairs found to locally co-occur (*n* = 441, see Methods), and accounting for the effects of divergence time, dispersal ability, and latitude, we found strong evidence that sympatry is associated with greater divergence in body mass, and, to a lesser extent, beak morphology (Fig. 2, Extended Data Fig. 1, Supplementary Tables 4-5, 8-9, 18-19; see Methods). The relationship between large body mass differences and increased likelihood of sympatry was highly consistent across sensitivity analyses (Fig. 2, Extended Data Fig. 1, Supplementary Tables 4-5, 8-9, 18-19). These results are largely in agreement with previous studies showing that the transition from secondary contact to sympatry is facilitated by divergence in body mass and beak morphology^10,12^, and further suggest that divergence in body mass is a more critical factor.

**Figure 2.**
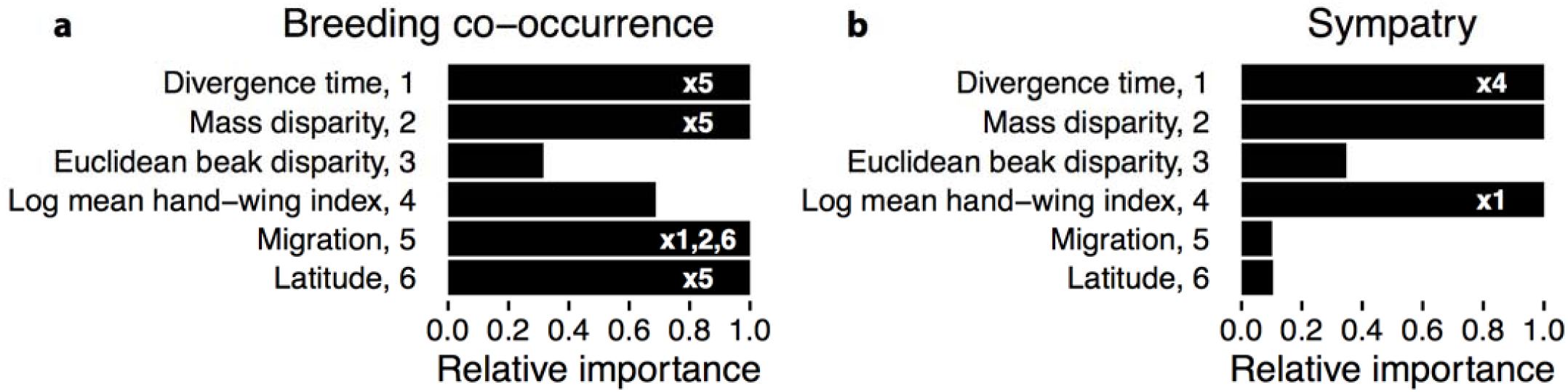
Factors associated with the establishment of secondary contact and sympatry in birds. Results of generalized linear models assessing the relative importance of predictors of breeding range local co-occurrence (**a**) and sympatry (**b**) in 952 avian sister species. Pairs with breeding co-occurrence include both parapatric and sympatric species pairs. Relative importance is estimated as the proportion of the summed model weights for all models with ?AIC<2, and indicates the extent to which each variable predicts the probability of co-occurrence or sympatry. Pairwise interactions with relative importance >0.6 are indicated by the numbers within the bar for each variable.

We have shown that secondary contact occurs earlier in the speciation process than generally assumed under classic models of allopatric speciation (Extended Data Figs 2, 4 and 7), suggesting a potentially wider role for parapatric speciation (speciation with no stage a in Fig. 1)^5^. Under this model of speciation, divergence pulses occur despite contact, and thus the potential for gene flow, between incipient species^15^. Our analyses indicate that this scenario may be widespread in bird speciation. However, we also found that the observed pattern of breeding co-occurrence and divergence times could also result from purely allopatric speciation with rapid rates of transition to secondary contact (speciation with reduced stage a in Fig. 1). Stochastic models indicate than an approximate minimum rate of 0.3 transitions to secondary contact per million years is sufficient to explain the pattern of breeding co-occurrences among avian sister species pairs (Extended Data Fig. 5, Supplementary Information). Thus, trait divergence pulses may take place either during periods of allopatry or parapatry, with cases of both probably widespread.

The standard evolutionary trajectory implied by our best-fitting model—an early, pulsed divergence with constrained subsequent divergence—can potentially explain a variety of phenomena in bird speciation. Under this single pulse model, pulses vary in magnitude across species, with a fraction of species undergoing large evolutionary jumps early in the speciation process, and others incurring small-magnitude divergence^17^. Thus, many sister species strongly resemble each other in ecological traits whether they began to diverge recently or anciently, whereas a fraction of species pairs have undergone an early pulse of rapid divergence, and remain highly divergent regardless of their age. One interpretation of this pattern is that a species pair undergoing a small pulse of ecological trait divergence in the early stage of speciation is unlikely to undergo large pulses at later stages in the absence of other speciation events, and thus the species pair may be subject to extended periods of mutual exclusion via ecological competition, perhaps in combination with reproductive interference^5,9^.

A prevailing view is that strong divergence in ecological traits between lineages typically requires long periods of time, i.e. slow-rate gradual divergence^12^. Gradual divergence models can account for prolonged mutual exclusion between highly similar species, a widespread phenomenon^5^, especially when evolutionary rates of gradual evolution are low^2,10^. However, gradual divergence models inadequately account for highly divergent young species pairs, unless they also incorporate brief bursts of faster gradual divergence (mimicking pulses)^27^. While gradual models thus may adequately explain avian trait evolution at macroevolutionary scales, our findings suggest that patterns of diversification among sister pairs are better captured by models incorporating pulses of divergence, where the magnitude of pulses is independent of divergence time. In particular, the single pulse model receives strong support and can help to explain the full range of outcomes observed in nature, which not only include prolonged mutual exclusion between ecologically similar sister species^6^ but also instances of rapid sympatry following abrupt ecological trait divergence^4^.

The occurrence of early pulses of trait divergence raises the question of how such pulses arise. One possibility is that they result from the intermittent discovery by populations of unoccupied adaptive peaks, as is expected in niche-filling models of diversification^8^?a mechanism that may be particularly important for large-magnitude pulses^4^. New adaptive peaks be discovered when local adaptation drives rapid divergence after range expansion^14,22^, for instance immediately following colonization of novel environments. Rapid phenotypic divergence may also result from local adaptation along environmental gradients, with or without gene flow^28^. In such contexts, it is worth emphasizing that signals of pulsed divergence may arise from a combination of gradual local (clinal) adaptation and subsequent extinction of intermediate populations^16,29^.

In combination, our results may help to resolve the longstanding question of why some nascent species survive over evolutionary time while others are ephemeral. One of the major threats to young lineages is the likelihood of fusion through swamping gene flow after secondary contact^16,21^. On one hand, we have shown this risk is widespread among nascent bird species because the lag time to secondary contact is shorter than expected (Extended Data Figs 2, 4-5), supporting the view that gene flow routinely becomes possible early in the speciation process. On the other hand, our findings suggest that species pairs undergoing major early pulses of ecological trait divergence are more likely to transition rapidly to sympatry, escaping both fusion and mutual exclusion, thereby extending their lifespan as independent lineages. Conversely, if they meet at early stages in the speciation process, species pairs with minimally divergent phenotypes may incur increased hybridization rates, or increased hybrid fitness, thereby reducing their lifespan. Indeed, elevated rates of extinction in less divergent young lineages may increase the signature of large early pulses in datasets compiled from extant species. Thus, differential extinction coupled with pulses of early trait divergence may play a critical role in explaining broad-scale patterns in the longevity and macroevolutionary diversity of species, as well as their geographical distributions.

## Acknowledgements

We are grateful to numerous data collectors who contributed to eBird, GenBank, and the CRC bird body mass data set (see Supplementary Information). We also thank Nico Alioravainen, Ed Braun, Samuel Jones, Rebecca Kimball, Dan Ksepka, Monte Neate-Clegg, Alex Pigot, Aaron Ragsdale and Gleb Zhelezov for data collection and technical assistance. This work was supported by the National Science Foundation (DEB-1208428 to J.G.B.), the Natural Environment Research Council (NE/I028068/1 to J.A.T.), and the Oxford Clarendon Fund and US-UK Fulbright Commission (to C.S.).

### Author contributions

J.G.B. and J.P.M. conceived the study; J.G.B, J.P.M. and J.A.T. designed the conceptual framework and analyses; J.G.B. performed dating analyses and assembled phylogenetic, occurrence, and body mass information; J.A.T. and C.S. provided morphometric data; J.P.M. integrated data sets, and designed and performed statistical analyses with significant input from J.G.B.; J.P.M. produced figures and tables; J.P.M. wrote the manuscript, with significant input from all authors.

### Author Information

Reprints and permissions information is available at www.nature.com/reprints. The authors declare no competing financial interests. Readers are welcome to comment on the online version of this article at www.nature.com/nature. Correspondence and requests for materials should be addressed to.

## METHODS

### Sister pairs

We used the maximum likelihood topology of Burleigh *et al*.’s^23^ avian supermatrix phylogenetic tree (hereafter “Burleigh tree”), which contains 6,714 species of the ∼10,500 bird species in the world, to select all (n = 2,076) pairs of avian sister species (i.e. each other’s closest relatives). The inclusion of some pairs of non-sister lineages would not invalidate our analyses but we tried to minimize this issue by excluding pairs that were unlikely to represent true sister species. Specifically, we excluded 763 pairs belonging to genera with <75% species-level sampling, and another 62 pairs that were deemed either unlikely to be true sister species based on molecular evidence from other studies or presented taxonomic problems. We further removed 299 species pairs for which we could not adequately score co-occurrence (Supplementary Dataset 6). This generated a manageable sample (n = 952) for downstream data quality checks. Different analyses use subsets of these 952 pairs depending on data availability and quality (see Supplementary Information Datasets 1-6).

### Divergence times

Divergence time estimates were obtained from a penalized likelihood analysis implemented in r8s^30^, using the maximum likelihood topology and molecular branch lengths from the Burleigh tree. We used 20 carefully-vetted fossil calibrations^24^ and constrained the root of the tree to a maximum age of 110 mya; applying an age constraint made little difference to estimated sister pair divergence times (see Extended Data Fig. 6). A list of the fossil calibrations (Supplementary Information Dataset 6) and a command block for the r8s analyses are available in the Supplementary Information. We performed sensitivity analyses using alternate sets of divergence time estimates, both from bootstrap analysis of the Burleigh tree, and from an independent phylogenetic and dating analysis^25^ (see Supplementary Information).

### Ecological trait measurements

*Body mass* Divergence in body size may be a strong contributor to ecological divergence, potentially reducing interspecific competition^4,31^ or reproductive interference^9^. We compiled data on body mass (a proxy for body size) from updated global datasets32-34. When multiple body mass values were reported, we took the mean; when male and female body masses were reported separately, we calculated an average of the two sex-specific means. We estimated body mass divergence as the difference between species in natural log of mean body mass^17^.

*Beak morphology* Species with similar body mass may partition niches according to diet. Thus, to quantify differences in foraging ecology among sister species, we collected three beak measurements (culmen length, beak depth, beak width) associated with food item selection and manipulation^10,35, 36^. Culmen length was measured as the distance from the distal part of the nostril to the beak tip. Beak depth and beak width were both measured at the distal edge of the nostril. All beak measurements were made on wild birds or museum specimens using calipers to the nearest 0.1mm (n ≥ 4 individuals sampled per species, 2 males and 2 females, where possible). See Supplementary Information for further details and rationale. To account for colinearity between beak measurements, we performed a phylogenetic Principal Components Analysis (phylogenetic PCA^37^; see Supplementary Table 1 for PC loadings).

*Dispersal* Highly vagile taxa with greater dispersal capacities should undergo faster range expansions, leading to earlier secondary contact in nascent species^38^. This can be associated with faster transition rates to sympatry^39^, but when secondary contact is very early, it may also slow or reverse the speciation process by promoting gene flow, leading to merged gene pools rather than coexistence^22^. Because of the importance of dispersal in allopatric speciation models, we assess how dispersal capacity influences transitions from allopatry to secondary contact, and from secondary contact to sympatry, respectively. As it is difficult to measure dispersal capacity directly, we instead used the hand-wing index (HWI), an index of wing shape related to the aspect ratio of the wing^38^ and a proxy for flight performance^40^. Using measurements (to the nearest mm) taken from wild birds and museum specimens, we calculated this index as

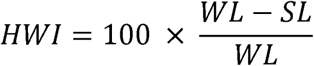

where WL (wing length) is the length of the closed wing from carpal joint to wing tip, and SL (secondary length) is the distance from the carpal joint to the tip of the first secondary feather. As a secondary index of dispersal, we also used range maps^41^ to assess migratory behaviour. If either member of a pair was illustrated as migratory to any degree, the species pair was scored as migratory.

### Geographical phases of speciation

*Secondary contact* We estimated local co-occurrence using ∼178 million bird species observation records stored in the eBird observational record database^42,43^. For a given species pair, local co-occurrence was defined as the occurrence of both species on the same day at the same reported locality. We also produced a narrower dataset of breeding range local co-occurrence by checking the dates and localities of co-occurrence records against breeding range maps^41^ and breeding phenology32. To qualify as evidence of breeding range local co-occurrence, species had to be reported on the same day and in the same locality during known breeding seasons of both species^32^ and within the known breeding range of one of the two species^41^. All sister species pairs for which breeding local co-occurrence has been documented were considered to have established secondary contact for the purposes of downstream analyses.

Because co-occurrence is unlikely to be reported for species with very few observations, we excluded sister pairs where at least one species had fewer than 10 eBird sightings reported. Our co-occurrence scores likely underestimate the true extent of co-occurrence among species pairs, as even after this filtering process, the minimum number of observations strongly predicts the probability of species pair local co-occurrence in our data set (GLM with the log of the minimum observations as sole predictor: coefficient estimate = 3.8 x 10^-4^ ± 8.5 x 10^-5^ SE; see Supplementary Information). Consequently, we conducted sensitivity analyses adopting minima of 20 and 50 observations (see Extended Data Fig. 9; Supplementary Tables 14-17). We also checked observational evidence for co-occurrence, discounting cases likely attributable to anthropogenic introductions and excluding cases potentially based on misidentifications or taxonomic confusion (see included and excluded species pairs in SI Datasets 1 and 6). *Sympatry* To examine the transition from secondary contact to wider coexistence (sympatry), we calculated percent breeding range overlap from geographic range polygons^41^ with a custom R script, using the R libraries rgdal, rgeos, maptools, and raster. A small subset of species pairs (n = 17 of 441 species pairs) could not be scored using our automated routine, and their range overlap was estimated visually. Species pairs with a range overlap >20% of the smaller range were scored as sympatric, while those with breeding co-occurrence but with ≤20% range overlap were scored as parapatric (having abutting ranges)^39^. Because some species pairs might best be considered sympatric even when their range overlap is less than 20%, we performed sensitivity analyses using an overlap of >10% scored as sympatric (Supplementary Tables 18-19).

### Analyses

#### Tempo of body mass divergence

To examine the tempo and timing of body mass and beak PC1 divergence, we investigated the relative support for four models of divergence for the species pairs from the full dataset for which body mass or beak morphology data were available (n = 869 species pairs for body mass, n = 926 species pairs for beak morphology). We fit one model of time-independent bounded evolution, and three different models that comprise a bounded evolution component on shorter timescales and one of three additional components for longer timescales^17^. These longer-timescale components are a gradual evolution model (Brownian motion) and two forms of pulsed divergence: a single pulse model where a single instantaneous displacement occurs following a waiting time sampled from an exponential distribution, and a multiple pulse model where the expected number of displacements for a given divergence time is determined by a Poisson process. We examined relative support for these models using AIC from likelihood calculations performed in R. We calculated confidence intervals for the Poisson rate parameter ?, the inverse of which is taken as the expected waiting time to a pulse, using likelihood profiling.

#### Secondary contact and sympatry

We examined the probability of local co-occurrence, and parapatry versus sympatry, using GLM with binomial error distributions, implemented in R^44^. In analyses of local co-occurrence and breeding range local co-occurrence for sister pairs, we began by predicting the probability of co-occurrence with divergence time as the only predictor (Extended Data Figs 3a-b and 7). We subsequently performed a model generation and selection routine (using the genetic algorithm of R package glmulti^45^, see Supplementary Information) to examine which among a set of phenotypic measures best predict local co-occurrence or sympatry while accounting for the effects of three variables that may influence the timing of transitions from allopatry to sympatry: divergence time^12^, latitude^9^ and dispersal ability^39^. The predictors of primary interest were between-species disparity in two traits implicated in ecological and reproductive isolation: body mass^46^ and beak morphology^47^. We incorporated disparity in beak morphology as (i) the Euclidean distance between species in PC space (following scaling of all PC’s to unit variance) or (ii) species differences along each PC axis as separate predictors. To account for differences among sister species pairs in dispersal ability, we also included the average log hand-wing index^38^ and migratory status of the sister pair as predictors. We further included divergence time and midpoint latitude (average of the two median observational latitudes for each species from eBird^43^). Our model generation routine permitted all pairwise interactions between predictors to enter the model, under the constraint that all models were marginal. We report support for all predictors entering the set of local co-occurrence models with ΔAIC < 2 (Supplementary Tables 2-3). All continuous variables were scaled and centered, such that estimated slope magnitudes for individual variables are meaningful in relation to one another.

For GLM examining the probability of sympatry versus parapatry, we first limited the sister species data set to those pairs that locally co-occur in breeding ranges. This restriction focuses the analysis on taxa that have the opportunity to interact to some degree in the breeding season^48^. The response variable in GLM is the geographic configuration: parapatric (interacting but without substantial range overlap) versus sympatric (having substantial range overlap: >20% of the smaller range in the analyses presented in the main text). We again used a genetic algorithm (see Supplementary Information) to generate model variants and performed model selection using the R package glmulti^45^.

To assess the sensitivity of our results to uncertainty in phylogenetic inference and divergence time estimates, we repeated all GLM analyses using mean divergence times for our species pairs from 100 samples of the pseudo-posterior distribution of trees from an alternative Bayesian species-level phylogenetic analysis^25^ (hereafter, the “Jetz tree”; Supplementary Information Tables S6-S9). For analyses examining the probability of local co-occurrence (and breeding local co-occurrence) with divergence time, we performed additional sensitivity analyses using divergence time estimates from 100 bootstraps of the Burleigh tree, and for each of the 10,000 psuedo-posterior samples from the Jetz tree.

#### Simulations of range dynamics

To aid in the interpretation of our GLM predicting local co-occurrence, we performed stochastic range dynamic simulations^39^. We used these simulations to place an approximate lower bound on the rate of secondary contact establishment from an initially allopatric configuration. To perform this estimation, we simulated the establishment of secondary contact using a simple model^10,39^, in which sister pairs can be in one of two states: co-occurring and not co-occurring. We simulated transitions into and out of contact over a set of possible rates from 0.1 to 0.8 per million years, in which the forward rate (rate of transition from isolation to contact, σ) is always greater than or equal to the reverse rate (rate of transition out of contact, ε). The forward and reverse rates are constant^39^, and the variation in rates among species arises only from stochasticity. Reverse rates were simulated at .005, .01, .05, .1, .2, and .5 times each of the forward rates. We present the maximum intercept calculated across all reverse rates (ε) for each simulated forward rate (σ) (Extended Data Figs 5, 12). To calculate the approximate percentage of species pairs coming into secondary contact by given points in time following divergence (100,000 years, 1 million years), we simulated range dynamics with σ = 0.3, and ε = .15 (corresponding to the minimal σ that yielded intercept >0.434, and the value of ε that yielded the highest intercept for σ = 0.3).

## References

1. Eldredge, N. & Gould, S. in Models in Paleobiology (Freeman, Cooper, San Francisco, 1972).

2. Pennell, M. W., Harmon, L. J. & Uyeda, J. C. Is there room for punctuated equilibrium in macroevolution? Trends Ecol. Evol. 29, 23-32 (2014).

3. Pfennig, D. W. & Pfennig, K. S. Character displacement and the origins of diversity. Am. Nat. 176 Suppl 1, S26–44 (2010).

4. Schluter, D. in The Ecology of Adaptive Radiation (Oxford University Press, Oxford, UK, 2000).

5. Price, T. in Speciation in Birds (Roberts and Company, Greenwood, Village, Colorado, 2008).

6. Rundell, R. J. & Price, T. D. Adaptive radiation, nonadaptive radiation, ecological speciation and nonecological speciation. Trends Ecol. Evol. 24, 394–399 (2009).

7. Mayr, E. in Systematics and the Origin of Species (Columbia University Press, 1942).

8. Price, T. D. et al. Niche filling slows the diversification of Himalayan songbirds. Nature 509, 222–225 (2014).

9. Weir, J. T. & Price, T. D. Limits to speciation inferred from times to secondary sympatry and ages of hybridizing species along a latitudinal gradient. Am. Nat. 177, 462–469 (2011).

10. Pigot, A. L. & Tobias, J. A. Species interactions constrain geographic range expansion over evolutionary time. Ecol. Lett. 16, 330–338 (2013).

11. MacArthur, R. & Levins, R. The limiting similarity, convergence, and divergence of coexisting species. Am. Nat., 377–385 (1967).

12. Tobias, J. A. et al. Species coexistence and the dynamics of phenotypic evolution in adaptive radiation. Nature 506, 359–363 (2014).

13. Smith, J. W. & Benkman, C. W. A coevolutionary arms race causes ecological speciation in crossbills. Am. Nat. 169, 455–465 (2007).

14. Friis, G., Aleixandre, P., Rodríguez-Estrella, R., Navarro-Sigüenza, A. G. & Milá, B. Rapid postglacial diversification and long-term stasis within the songbird genus Junco: phylogeographic and phylogenomic evidence. Mol. Ecol. 25, 6175–6195 (2016).

15. Rundle, H. D. & Nosil, P. Ecological speciation. Ecol. Lett. 8, 336–352 (2005).

16. Futuyma, D. J. On the role of species in anagenesis. Am. Nat. 130, 465–473 (1987).

17. Uyeda, J. C., Hansen, T. F., Arnold, S. J. & Pienaar, J. The million-year wait for macroevolutionary bursts. Proc. Natl. Acad. Sci. U. S. A. 108, 15908–15913 (2011).

18. Estes, S. & Arnold, S. J. Resolving the paradox of stasis: Models with stabilizing selection explain evolutionary divergence on all timescales. Am. Nat. 169, 227–244 (2007).

19. Bokma, F. Time, species, and separating their effects on trait variance in clades. Syst. Biol. 59, 602–607 (2010).

20. Rabosky, D. L. & Adams, D. C. Rates of morphological evolution are correlated with species richness in salamanders. Evolution 66, 1807–1818 (2012).

21. Futuyma, D. J. in Macroevolution 29-85 (Springer, 2015).

22. Mayr, E. in Animal Species and Evolution (Belknap Press of Harvard University Press, Cambridge, Massachusetts, 1963).

23. Burleigh, J. G., Kimball, R. T. & Braun, E. L. Building the avian tree of life using a large-scale, sparse supermatrix. Mol. Phylogenet. Evol. 84, 53–63 (2015).

24. Baiser, B., Valle, D., Zelazny, Z. & Burleigh, J. G. Non-random patterns of invasion and extinction reduce phylogenetic diversity in island bird assemblages. Ecography (2017).

25. Jetz, W., Thomas, G. H., Joy, J. B., Hartmann, K. & Mooers, A. O. The global diversity of birds in space and time. Nature 491, 444–448 (2012).

26. Arnold, S. J. Phenotypic evolution: the ongoing synthesis (American Society of Naturalists address). Am. Nat. 183, 729–746 (2014).

27. Landis, M. J., Schraiber, J. G. & Liang, M. Phylogenetic analysis using Levy processes: finding jumps in the evolution of continuous traits. Syst. Biol. 62, 193–204 (2013).

28. Endler, J. A. in Geographic Variation, Speciation, and Clines (Vol. 10) (Princeton University Press, 1977).

29. Futuyma, D. J. Evolutionary constraint and ecological consequences. Evolution 64, 1865–1884 (2010).

30. Sanderson, M. R8s: Inferring absolute rates of molecular evolution and divergence times in the absence of a molecular clock. Bioinformatics 19, 301–302 (2003).

31. Wilson, D. S. The adequacy of body size as a niche difference. Am. Nat. 109, 769–784 (1975).

32. del Hoyo et al. (eds.). Handbook of the Birds of the World Alive. Lynx Edicions, Barcelona. (retrieved from http://www.hbw.com/ in 2015-16).

33. Dunning, J. B. in Body Masses of Birds of the World (Taylor and Francis Group, Boca Raton, Florida, 2008).

34. Dunning, J.B. Body Masses of Birds of the World. http://ag.purdue.edu/fnr/Documents/WeightBookUpdate.pdf. (2016).

35. Schoener, T. W. The evolution of bill size differences among sympatric congeneric species of birds. Evolution 19, 189–213 (1965).

36. Miles, D. B. & Ricklefs, R. E. The correlation between ecology and morphology in deciduous forest passerine birds. Ecology 65, 1629–1640 (1984).

37. Revell, L. J. phytools: an R package for phylogenetic comparative biology (and other things). Methods Ecol. Evol. 3, 217–223 (2012).

38. Claramunt, S., Derryberry, E. P., Remsen, J. V.,Jr. & Brumfield, R. T. High dispersal ability inhibits speciation in a continental radiation of passerine birds. Proc. R. Soc. Lond. B 279, 1567–1574 (2012).

39. Pigot, A. L. & Tobias, J. A. Dispersal and the transition to sympatry in vertebrates. Proc. R. Soc. Lond. B 282, 20141929 (2015).

40. Dawideit, B. A., Phillimore, A. B., Laube, I., Leisler, B. & Böhning-Gaese, K. Ecomorphological predictors of natal dispersal distances in birds. J. Anim. Ecol. 78, 388–395 (2009).

41. BirdLife International and NatureServe. in Bird species distribution maps of the world (BirdLife International and Natureserve, Cambridge, UK and Arlington, USA, 2014).

42. Sullivan, B. L. et al. eBird: A citizen-based bird observation network in the biological sciences. Biol. Conserv. 142, 2282–2292 (2009).

43. Sullivan, B. L. et al. The eBird enterprise: An integrated approach to development and application of citizen science. Biol. Conserv. 169, 31–40 (2014).

44. R Core Team. R: A language and environment for statistical computing. (2012).

45. Calcagno, V. & de Mazancourt, C. glmulti: an R package for easy automated model selection with (generalized) linear models. J. Stat. Softw. 34, 1–29 (2010).

46. Yasukawa, K. Male quality and female choice of mate in the red-winged blackbird (Agelaius phoeniceus). Ecology 62, 922–929 (1981).

47. Grant, P. R. & Grant, B. R. Hybridization, sexual imprinting, and mate choice. Am. Nat. 149, 1–28 (1997).

48. Lovette, I. J. & Hochachka, W. M. Simultaneous effects of phylogenetic niche conservatism and competition on avian community structure. Ecology 87, S14–S28 (2006).

